# Unbiased dynamic characterization of RNA-protein interactions by OOPS

**DOI:** 10.1101/333336

**Authors:** Rayner M. L. Queiroz, Tom Smith, Eneko Villanueva, Mie Monti, Mariavittoria Pizzinga, Maria Marti-Solano, Dan-Mircea Mirea, Manasa Ramakrishna, Robert F. Harvey, Veronica Dezi, Sven Degroeve, Lennart Martens, Gavin H. Thomas, Anne E. Willis, Kathryn S. Lilley

## Abstract

Current methods for the identification of RNA–protein interactions require a quantity and quality of sample that hinders their application, especially for dynamic biological systems or when sample material is limiting. Here, we present a new approach to enrich RNA-Binding Proteins (RBPs): Orthogonal Organic Phase Separation (OOPS), which is compatible with downstream proteomics and RNA sequencing. OOPS enables recovery of RBPs and free protein, or protein-bound RNA and free RNA, from a single sample in an unbiased manner. By applying OOPS to human cell lines, we extract the majority of known RBPs, and importantly identify additional novel RBPs, including those from previously under-represented cellular compartments. The high yield and unbiased nature of OOPS facilitates its application in both dynamic and inaccessible systems. Thus, we have identified changes in RNA-protein interactions in mammalian cells following nocodazole cell-cycle arrest, and defined the first bacterial RNA-interactome. Overall, OOPS provides an easy-to-use and flexible technique that opens new opportunities to characterize RNA-protein interactions and explore their dynamic behaviour.

## Introduction

Interactions between RNA-binding proteins (RBPs) and RNA are essential for the control of gene expression by regulating the processes of transcription, trafficking, decay and mRNA translation^1–7^. In this way, RNA-RBP crosstalk controls cell homeostasis and cell fate. Several approaches have been developed to characterize this crosstalk. Studies of Protein-Bound RNA (PBR) generally require the purification, e.g. by immunoprecipitation, of a specific protein and subsequent sequencing of its RNA cargo ^8,9^. From the protein perspective, the repertoire of RBPs can be examined by UV crosslinking followed by capture of polyadenylated RNAs using oligo(dT) and the identification of the bound proteins^10–12^. However, despite their considerable contribution to the understanding of RNA-protein crosstalk, such methods have limitations. Thus, techniques designed to study PBRs are incompatible with a system-wide analysis of RBPs and PBRs, whereas oligo(dT)-based purification requires a large amount of starting material, with high reagent costs, which complicates its application in dynamic conditions^13^. Moreover, the theoretical dependence on the presence of RNA polyA tails, makes oligo(dT)-based capture methods incompatible for organisms with little or no polyadenylation, such as prokaryotes, or for non-polyadenylated RNAs; thus introducing bias in the RBPs identified. There have been recent attempts to address the oligo(dT) limitations, but these have involved incorporation of modified nucleotides, which by itself restricts its applicability and introduces a bias in transcription-dependent nucleoside-incorporation ^14–16^.

To address these limitations, we have developed a method based on Acidic Guanidinium Thiocyanate-Phenol-Chloroform (AGPC) phase partition, called Orthogonal Organic Phase Separation (OOPS). AGPC purification permits the recovery of RNA species in an unbiased manner^17,18^. In standard conditions, when lysing cells in AGPC, two distinct phases are formed: RNA migrates to the upper aqueous phase and proteins occupy the lower organic phase. Here, we utilise UV crosslinking at 254 nm to generate RNA-protein adducts that combine the physicochemical properties of both molecules and thus migrate to the aqueous-organic interface^19^. Isolation of the interface allows specific recovery of either RBPs or PBRs by digesting the reciprocal component of the adduct.

Here, we demonstrate the specificity and versatility of OOPS. Separation of free and protein-bound RNA provides a way to quantify the proportion of RNA crosslinked to protein, enabling precise optimisation of UV dosage. Furthermore, we show that OOPS recovers all crosslinked-RNA (CL-RNA) and thus all crosslinked RBPs. To demonstrate the versatility of OOPS, we analyse RNA-binding changes through cell-cycle progression following nocodazole arrest. Finally, we apply OOPS to describe the first bacterium RNA-interactome, confirming that OOPS is fully applicable for organisms with limited polyadenylation.

## Results

### OOPS specifically recovers protein-bound RNA

Cell lysis in acidic Guanidinium-Thiocyanate-Phenol, and subsequent addition of chloroform to the sample, generates two distinct phases: an upper aqueous phase containing RNA and a lower organic phase containing proteins. We reasoned that stable RNA-protein adducts, induced by UV-crosslinking at 254 nm, should therefore be retained at the interface (First step of Figure 1a). CL-RNA was recovered from the interface by protein digestion using proteinase-K and extracting the resulting RNA from the aqueous phase of a subsequent phase separation (Figures 1a-b, online methods). RNA migration from the interface to the aqueous phase after protein digestion indicates that its presence at the interface was dependent on protein binding. Importantly, we observed a dose-dependent depletion of RNA from the aqueous phase (Figures 1b, S1a) with CL-RNA saturating at approximately 75% (Figure 1b). The size profile of CL-RNA resembles total free-RNA of a non-crosslinked sample (NC), with the aqueous phase of the CL sample containing free small RNAs (Figure S1b).

**Figure 1.**
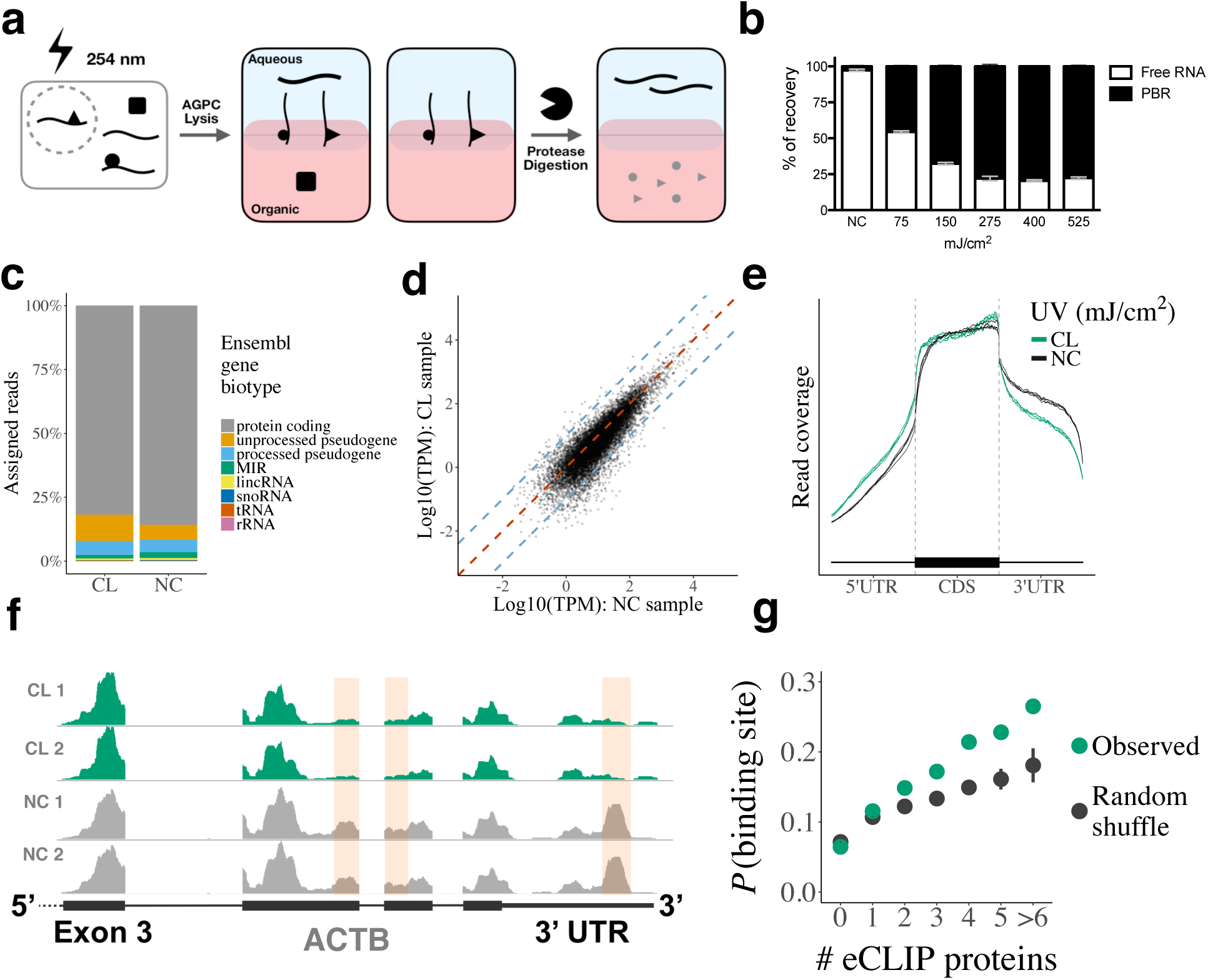
OOPS unbiasedly recovers all PBR species. **(a)** Schematic representation of the OOPS method to extract protein-bound RNA **(b)** Relative proportions of free RNA (aqueous phase) and protein-bound RNA (PBR; interface) with increasing UV dosage. Data shown as mean +/− SD of 3 independent experiments **(c)** Relative proportions of reads assigned to Ensembl gene biotypes for 400 mJ/cm^2^ CL and NC samples **(d)** Correlation between NC replicate 1 and 400 mJ/cm^2^ CL replicate 1. Blue dashed lines represent a 10-fold difference **(c)** Meta-plot of read coverage over protein-coding gene-model **(f)** Read coverage across the 3’ end of ATCB for CL (400 mJ/cm^2^) and NC replicates. Red boxes denote regions with consistently reduced coverage in CL **(g)** Relationship between the number of eCLIP proteins with a peak in a sliding window and the probability of the window being identified as a protein binding site Non-crosslinked=NC, Crosslinked=CL

To compare total RNA from a non-crosslinked sample and CL-RNA, rRNA was depleted from both conditions and total RNA-seq carried out, using a dose range of 150-400 mJ/cm^2^ (Figure S1c). The abundance of RNA species observed in the CL-RNA and NC-RNA samples was similar, with protein-coding RNAs predominating (Figures 1c, S1d). Crucially, the Pearson correlation between CL and NC samples is as high as the correlation among crosslinked samples (median correlations are 0.89 and 0.92, respectively; Figures 1d, S1e), suggesting CL-RNAs fully represent the long RNAs present in a cell and thus, all long RNA species are bound by protein.

Despite the high correlation between RNA abundance in CL and NC samples, we observed an overall reduction of coverage in the 3’ UTRs of mRNAs (Figures 1e) and a loss of coverage at discrete positions (Figure 1f). We hypothesized that this was due to steric hindrance of reverse transcription at sites of RNA-protein crosslinking, as protein binding to RNA occurs frequently within the 3’ UTR^20^. To test this, we applied a sliding window approach to identify the sites across the transcriptome where a significant loss of coverage was observed in CL samples (see supplementary notes). As expected, these sites were more frequent in mRNA 3’ UTRs and showed a significant overlap with ENCODE eCLIP protein-binding peaks^21^, confirming they represent positions of protein binding (Figure 1g, S1f). Windows with a higher uracil content, which preferentially crosslink to proteins at 254 nm, are more likely to contain a protein-binding site, but adjacent uridines, which are liable to photo-dimerize with 254 nm UV and block reverse transcription^22^, have no effect (Figure S1g). This further suggests that protein binding is the predominant cause of observed differences in read coverage.

### OOPS recovers RNA-binding proteins

Having established that OOPS allows efficient extraction of PBRs, we next identified the proteins crosslinked to RNA. Notably, this required less than 1% of the cells needed in the more established RBP-capture methods ^10,23^ (see online methods). First, we confirmed that the presence of proteins at the interface was dependent on UV-crosslinking. Stable Isotope Labeling by Amino acids in cell Culture (SILAC)^24^ quantitation was used to determine the relative abundance of proteins from CL and NC U2OS cells in the same OOPS interface (Figure 2a; see online methods). Importantly, repeated phase separation improves the protein CL:NC ratio of RBPs, indicating that additional phase separations remove non-crosslinked proteins. The 3rd interface appears to be optimal, as subsequent phase separations reduce the number of identified proteins (Figures 2b, S2a, supplementary table 1). As expected, since glycosylated proteins share the physicochemical properties of RNA-protein adducts, their presence at the interface is CL-independent. In contrast, non-glycosylated proteins show a similar CL-enrichment, regardless of their GO annotation as RBPs (Figures 2b). This confirms crosslinking enriches known RBPs, and potential undiscovered RBPs, in the interface.

**Figure 2.**
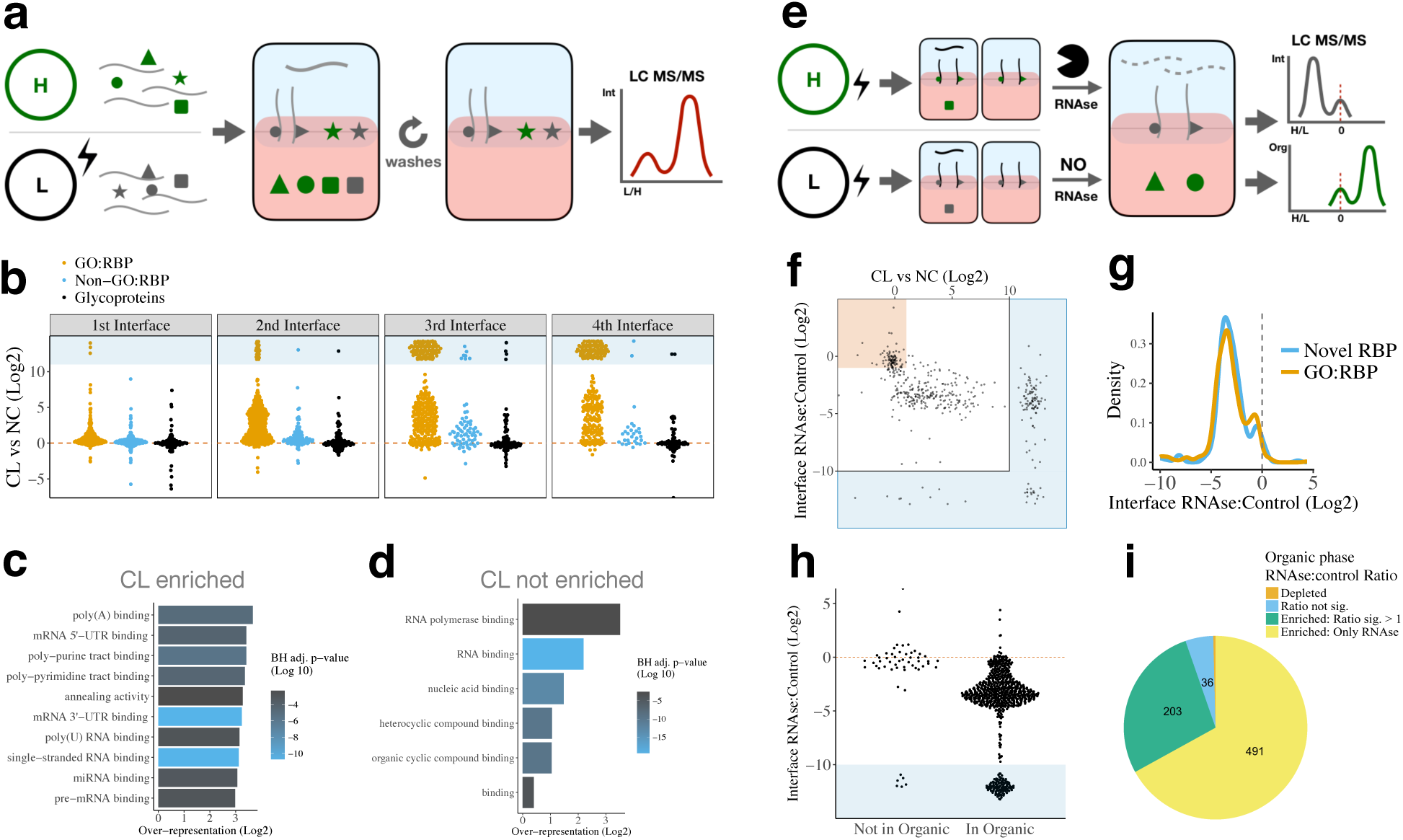
OOPS specifically recovers RBPs. **(a)** Schematic representation of the SILAC experiment used to determine the effect of UV crosslinking on protein abundance in the interface and the effect of additional phase separation cycles to wash the interface **(b)** Protein CL vs NC ratios for the 1^st^ to 4^th^ interfaces **(c)** Top 10 molecular function GO terms over-represented in proteins enriched by CL in the 3^rd^ interface **(d)** Top 10 molecular function GO terms over-represented in proteins not enriched by CL in the 3^rd^ interface **(e)** Schematic representation of a SILAC experiment to determine protein abundance in the 3rd interface and 4th organic phase following RNAse treatment **(f)** Protein CL vs NC ratio and RNAse vs control ratio in the interface. Red box denotes proteins which are not CL-enriched and not depleted by RNAse. The regions around the graph in blue denote ratios which cannot be accurately estimated as they are > 2^10^ or < 2^−10^, or the protein was only detected in one condition **(g)** RNAse vs control ratio in the interface for GO annotated RBPs and other OOPS RBPs **(h)**Protein RNAse vs control ratio in the RNAse treated interfaces for proteins identified in the 4th step organic phase **(i)** Proportion of proteins enriched in the organic phase following RNAse treatment

Excluding glycoproteins, 70% of proteins were enriched at the 3rd interface post UV-crosslinking (Figure S**2b-e**). A similar proportion of proteins were enriched with a lower dosage (150 mJ/cm^2^; Figure S2e). CL-enriched proteins showed a clear over-representation of RNA-related GO terms (Figure 2c). Remarkably, even within the CL-independent proteins, and accounting for protein abundance, there was a clear over-representation of RNA-binding GO terms (Figure 2d), suggesting that CL-enrichment alone is not sufficient to distinguish free proteins from RNA-bound proteins.

In order to establish that the presence of the proteins at the interface was RNA-dependent, we treated the interfaces with ribonucleases (RNase), and determined protein migration to the organic phase (Figure 2e; online methods, Figures 2f, S2f). Notably, the proteins which migrated to the organic phase included proteins which were CL-independent, suggesting their presence in the interface was RNA-dependent (Figure 2f, S2i). Moreover, proteins not annotated as RBPs show similar RNase sensitivity to those annotated as RBPs, suggesting these are undiscovered RBPs (Figure 2g). In contrast, glycoprotein abundance at the interface was not affected by RNase (Figure S2g). Since the presence of glycoproteins at the interface was also CL-independent (Figure 2b), we excluded them from downstream analyses. Ninety-three percent of proteins observed in the organic phase were RNase sensitive according to their reduction in interface abundances, whereas those missing from the organic phase were largely RNAse insensitive (Figure 2h). As expected, 95% of proteins extracted from the organic phase showed an enrichment following RNAse treatment (Figure 2i). These proteins show a clear over-representation of GO terms related to RNA binding (Figure S2i). Together, these experiments, demonstrate that RNAse treatment is a necessary step to ensure extraction of *bona fide* RBPs. Importantly, these data were replicated in HEK293 cells, confirming cell line-independence (Figure S2c-h).

### OOPS identifies canonical and novel RBPs in different cellular compartments

The RBPs identified by OOPS were compared to those from oligo(dT) RBP-Capture analysis. We confirmed a high degree of overlap, with 78% of proteins identified by RBP-capture in U2OS cells also identified by OOPS (Figure 3a, S3a for HEK293 cells). Furthermore, even amongst the proteins identified by only one method, there was a significant over-representation of GO-annotated RBPs (p-value < 2.2e-16, Fisher’s Exact Test). We next applied OOPS to identify RBPs from MCF10A (a cell line derived from a healthy individual), and observed a “common” RBPome of 619 proteins detected in all 3 cell lines (Figure 3b, S3b, supplementary table 2) and a minority of cell line-specific RBPs. A comparison of the 1555 of the proteins from the 3 cell lines used in this study with all previous human RBP-capture data, showed 70% identity (Figure 3c). In addition, OOPS identified 80% of the proteins isolated by polyA-independent RICK^15^ and CARIC^14^ methods (Figure S3c-d). These results indicate that OOPS is able to recover the majority of the annotated RBPome, including proteins that do not bind poly-adenylated RNAs.

As expected, OOPS proteins show a clear over-representation of GO terms describing all forms of RNA-binding, including 5’ and 3’ UTR binding, and single and double-stranded RNA-binding (Figure 3d, S3e-f)). Interestingly, the novel RBPs identified by OOPS show an over-representation of GO terms related to mRNA transport and RNA localisation (Figure 3e, S3g). We therefore projected OOPS proteins onto our previous hyperLOPIT data^25^, which identifies the average localisation of proteins, as an initial indication of the subcellular localisation of the RNA-bound fraction. Known RBPs are predominantly localised to the nucleus, mitochondria, cytosol and large protein complexes (e.g ribosomes; Figure 3f), whereas OOPS RBPs appear to be more broadly distributed with a greater proportion of membrane proteins and proteins of unknown localisation (Figure 3f). Since membrane proteins are generally underrepresented in mass spectrometry experiments, we performed a crude cell fractionation to separate cellular compartments into 3 fractions: “heavy membranes” (e.g. nucleus, mitochondria), “light membranes” (e.g. ER, PM, etc.) and “cytosol” (Figure S3h). RBPs were detected from all fractions with the membrane fractions yielding more novel membrane-RBPs (Figures 3f, S3h-I, supplementary table 3). Moreover, we confirmed that transmembrane-domain containing RBPs were more abundant in the membrane fractions (Figure S3i). Taken together, our data indicate that combining OOPS with fractionation allows recovery of novel RBPs from previously underrepresented compartments.

**Figure 3.**
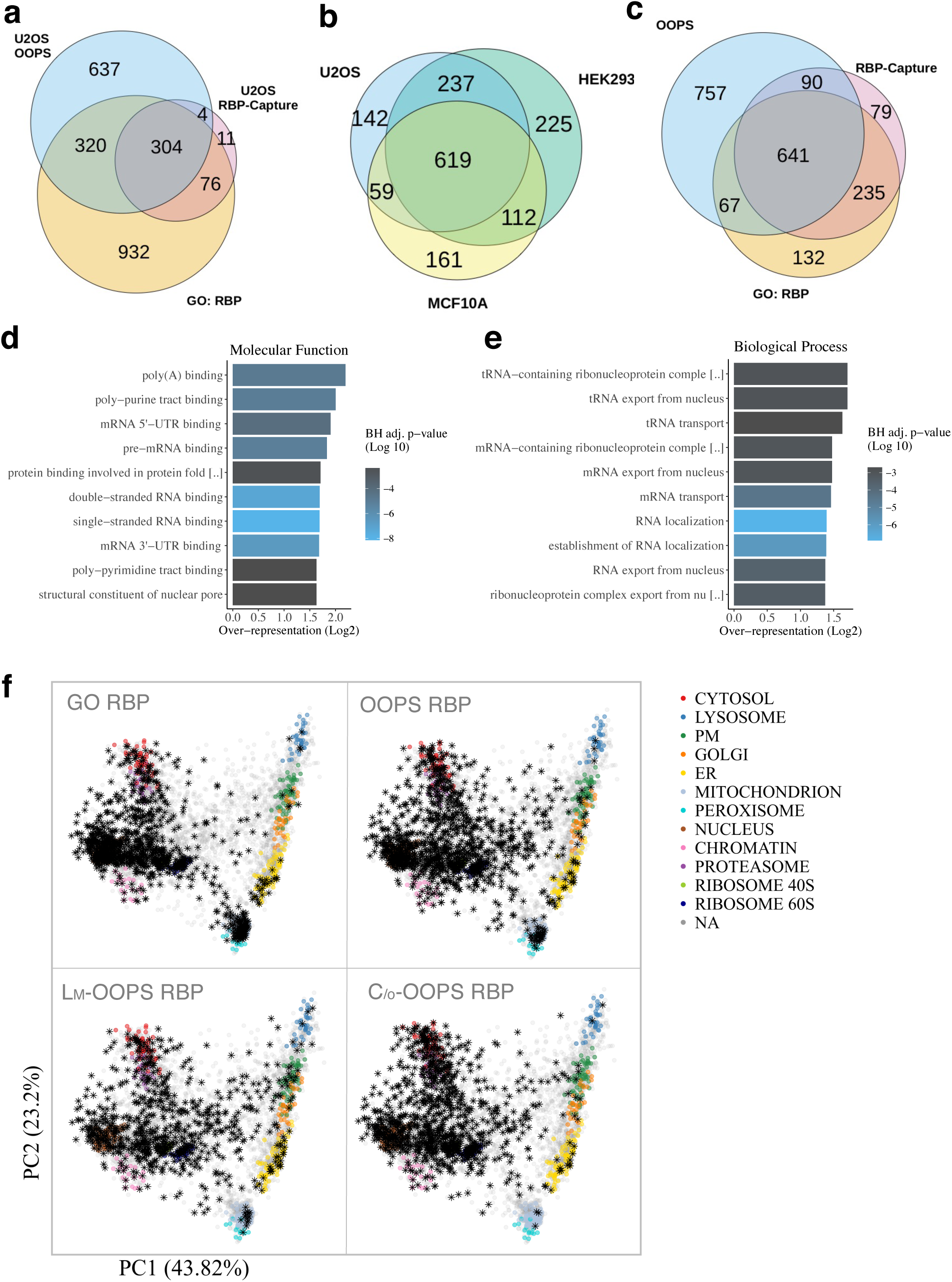
OOPS recovers known and new RBPs, even from under-represented cell compartments. **(a)** Overlap between OOPS, RBP-Capture and GO-annotated proteins for U2OS cells. Proteins were restricted to those expressed in U2OS **(b)** Overlap between proteins identified with OOPS from U2OS, HEK293 and MCF10A. Proteins were restricted to those expressed in all cell lines **(c)** Overlap between the union of OOPS proteins identified in the 3 cell lines, all published RBP-Capture studies, and GO annotated RBPs. Proteins were restricted to those expressed in at least one of the three OOPS cell lines **(d)** Top 10 molecular function GO terms over-represented in the proteins identified in U2OS, HEK293 or MCF10A OOPS **(e)** Top 10 biological process GO terms over-represented in novel RBPs identified by OOPS **(f)** HyperLOPIT projections of protein localisation. Canonical subcellular localisation markers indicated in colour as shown. NA: non-marker protein. Highlighted RBPs shown as black asterisks. Light membrane-enriched fraction: Lm, Cytoplasm/Other fraction: C/o

### High-throughput identification of the RNA-binding sites confirms OOPS RBPs

To validate our novel RBPs and further characterise their RNA binding sites, we developed a method to identify the precise RNA-binding sites, inspired by RBD-map^23^. Briefly, proteins from interfaces were sequentially digested to yield the peptide crosslinked to RNA (enriched by TiO_2_) and the peptides adjacent to the crosslinked site (See online methods; Figure 4a). Importantly, the RNA-peptide enrichment techniques used were orthogonal to OOPS, providing independent validation as well as yielding RNA-binding sites. Overall, we identified discrete putative RNA-binding sites in 578 (46%) of OOPS U2OS proteins using the adjacent peptides. As expected, putative binding sites were more easily identified in proteins with a higher abundance in the interface, with a binding site identified for 57% of the most abundant novel RBPs (Figure 4b, supplementary table 4). Direct determination of the crosslinked peptide using the novel Ionbot search engine (see supplementary notes), confirmed a significant overlap between the two approaches (Fig S4a).

**Figure 4.**
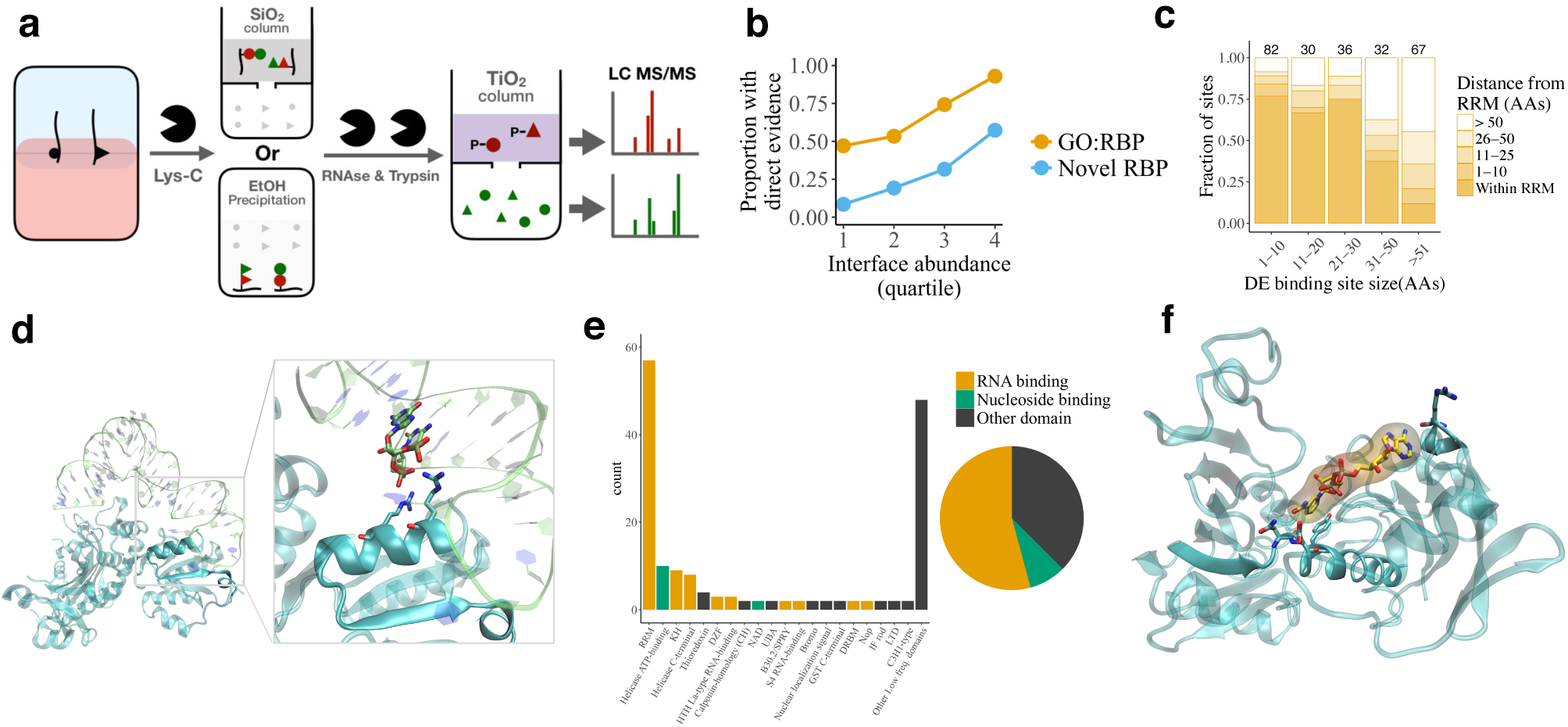
Direct assessment of crosslink sites validates most OOPS RBPs. **(a)** Schematic representation of the sequential digestion method used to identify the RNA-binding site. Red=peptides containing site of crosslinking. Green=peptides adjacent to the RNA-binding site peptide **(b)** Proportion of OOPS proteins in which the RNA-binding site was identified. Proteins separated into GO annotated RBPs and novel RBPs, and by their abundance at the OOPS interface **(c)** Distance of putative RNA-binding sites to the nearest RRM. Smaller putative RNA-binding sites are closer to RRM. Analysis restricted to proteins with an RRM **(d)** Crystal structure of Glycyl-tRNA synthetase in complex with tRNA-Gly (PDB ID 4KR2). RNA is shown as transparent lime ribbon, while the Glycyl-tRNA synthetase is shown in a cyan transparent cartoon representation. The protein region detected by direct evidence is shown in an opaque representation and RNA and protein residues at 4 Å or less from each other are shown as lime and cyan sticks respectively **(e)** Domains containing a putative RNA-binding site **(f)** Crystal structure of GAPDH complexed with NAD (PDB ID 4WNC). GAPDH is shown as a cyan transparent cartoon and the protein region detected by direct evidence is shown in an opaque representation. Residues at 4 Å or less from NAD (yellow sticks and surface representation) are shown as cyan sticks

To confirm the specificity of our approach, we focused on proteins containing an annotated RNA-recognition motif (RRM), and observed a significant overlap between the sites identified and RRMs (Figure 4c). To further test the precision of our RNA-binding site assignment, we examined crystal structures of our detected RBPs in complex with RNA. As an example, the crystal structure of the glycyl-tRNA synthetase in complex with tRNA-Gly^26^ confirms that the detected binding site contains residues less than 4 Å to the crystallized tRNA (Figure 4d). Additionally, we identified RNA-binding sites in 17 proteins in the ribosome quality control complex structure, together with a novel RBP whose binding site is also within 4 Å of the ribosomal RNA^27^ (Fig S4b). Finally, we established that our method identifies known RNA-binding domains in GO annotated RBPs, including the canonical RRM and KH domains, and non-canonical helicase C-terminal^28,29^ and DZF^23^ domains (Figure 4e). Alongside these non-canonical RNA-binding domains, we identified multiple NAD-binding domains (Figure 4e). These included two sites within the NAD-binding pocket of GAPDH^30^, which confirmed previous RNA-binding site predictions based on *in vitro* experiments^31^ (Figure 4f).

### OOPS facilitates quantification of RNA-binding in a dynamic system

To demonstrate the high yield and versatility of OOPS, we applied it to a dynamic system. U2OS cells were arrested in prometaphase (M) with nocodazole and dynamic changes in RNA-binding were determined following a short (6 h; G1) and long recovery (23 h; de-synchronized cells; using TMT quantification (Figures 5a & b, S5a and online methods). As expected, we observed changes in the abundance of cell-cycle related proteins between M and the post-release phases (Figure S5b). Quantifying protein abundance in OOPS and total cell lysates from the same sample (Figure 5b), enabled us to detect changes in RNA-binding, which were distinct from concurrent changes in total protein abundance. Interestingly, the changes in OOPS-enriched protein abundance frequently did not correlate with variations in total protein abundance, suggesting specific RBPs bind RNA differentially across the different cell-cycle stages (Figure 5c, supplementary table 5).

**Figure 5.**
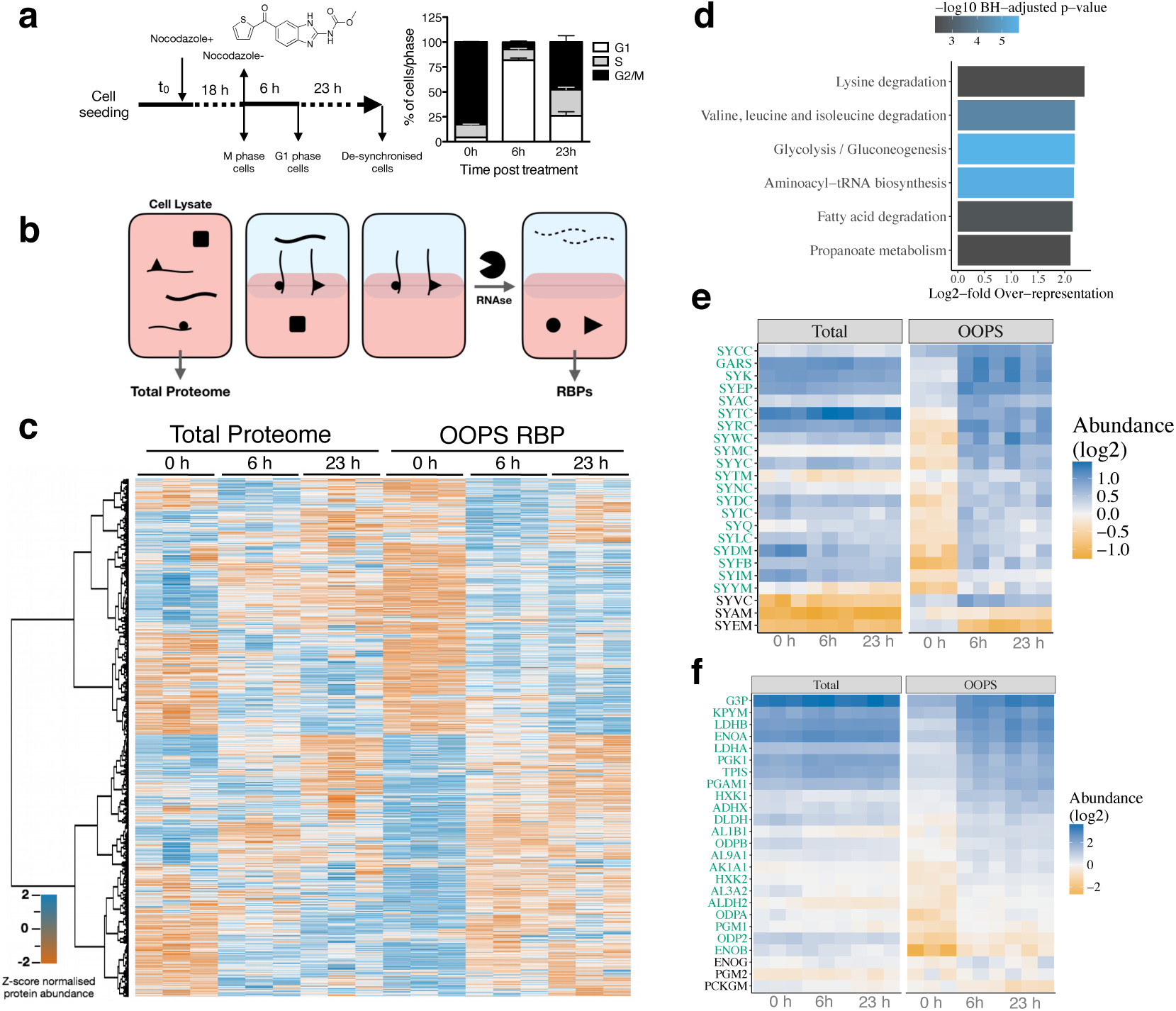
Dynamic characterization of the RBP-ome after nocodazole arrest. **(a)** Left: schematic representation of the nocodazole arrest/release experiment. Cells were analysed after 18 h nocodazole arrest and after a 6 h or 23 h release from the treatment release. Right: relative proportions of cells in G1, S and M phase for cells synchronised at each time-point (shown as the mean +/− SD of 3 independent experiments) **(b)** Schematic representation of protein extraction for nocodazole-arrest experiment. Total proteomes were extracted from cell lysates and RBPs were extracted following OOPS proteome method **(c)** Protein abundance from total proteome and OOPS extractions. Abundance z-score normalised within each extraction type. Proteins hierarchically clustered across all samples as shown on left **(d)** KEGG pathways over-represented in the RBPs with a significant increase in RNA-binding (**e-f**) Protein abundance for KEGG pathways over-represented in proteins with a significant increase in RNA-binding. Individual proteins with a significant increase are highlighted in green. **(e)** tRNA aminoacylation pathway. (**f**) Glycolysis pathway

To better understand the dynamics of these proteins, we used a linear model framework to identify proteins with apparent changes in RNA-binding, taking into account their total abundance (see online methods). We focused on changes occurring between arrested cells and 6 h release since these were the most synchronised time points. KEGG-pathway^32^ over-representation analysis indicates that many of the proteins that showed dynamic RNA-binding between these stages were involved in metabolism (Figure 5d). Interestingly, 20/23 tRNA synthetases detected showed an increase in RNA binding post nocodazole release, suggesting a coordinated increase in aminoacyl-tRNA availability (Figure 5e). In addition, we observed an over-representation of proteins involved in glycolysis (Fig5d). Importantly, the presence of glycolysis related proteins in OOPS interfaces was CL-dependent and RNase sensitive in our previous SILAC experiments (Figure S5c). Many of the proteins that function in glycolysis have been described as eukaryotic RNA-binding proteins previously ^12,33,34^. However, this is the first demonstration of dynamic RNA-binding for these RBPs (Figure 5f), providing a potential model to study the RNA-binding function of these proteins.

### OOPS reveals the first RBPome of a non-polyadenylated organism

Since OOPS is not restricted to the study of polyA-RNA binding proteins, we used this method to explore the RBPome of *E. coli* (see online methods for OOPS in bacteria). We detected 397 proteins, which represents 9% of the predicted proteome for the studied K-12 strains^35^, and is approximately equivalent to the proportion obtained in eukaryotic cells. We recovered 89/172 GO annotated RBPs (Figure 6a, supplementary table 2) and observed that the over-represented GO terms for OOPS proteins are related with the full repertoire of RNA biotypes, including the terms “tRNA aminoacylation”, “ribosome assembly” and “ribonucleoprotein complex assembly” (Figure 6b). As expected, the most enriched protein domains are related to nucleic acid binding (Figure 6c). In addition, helicase and P-loop domains, previously described in protein-RNA interactions are also present ^28,29,36^. Furthermore, if we only consider the 308 novel OOPS-RBPs, we still find a clear enrichment of RNA-assoicated GO-terms, the great majority of them related to tRNAs or ncRNAs (Figure 6d). However, 259 OOPS-RBPs are not annotated with an RNA-related GO term, suggesting OOPS provides a useful approach to study new RBP functions in prokaryotic systems.

**Figure 6.**
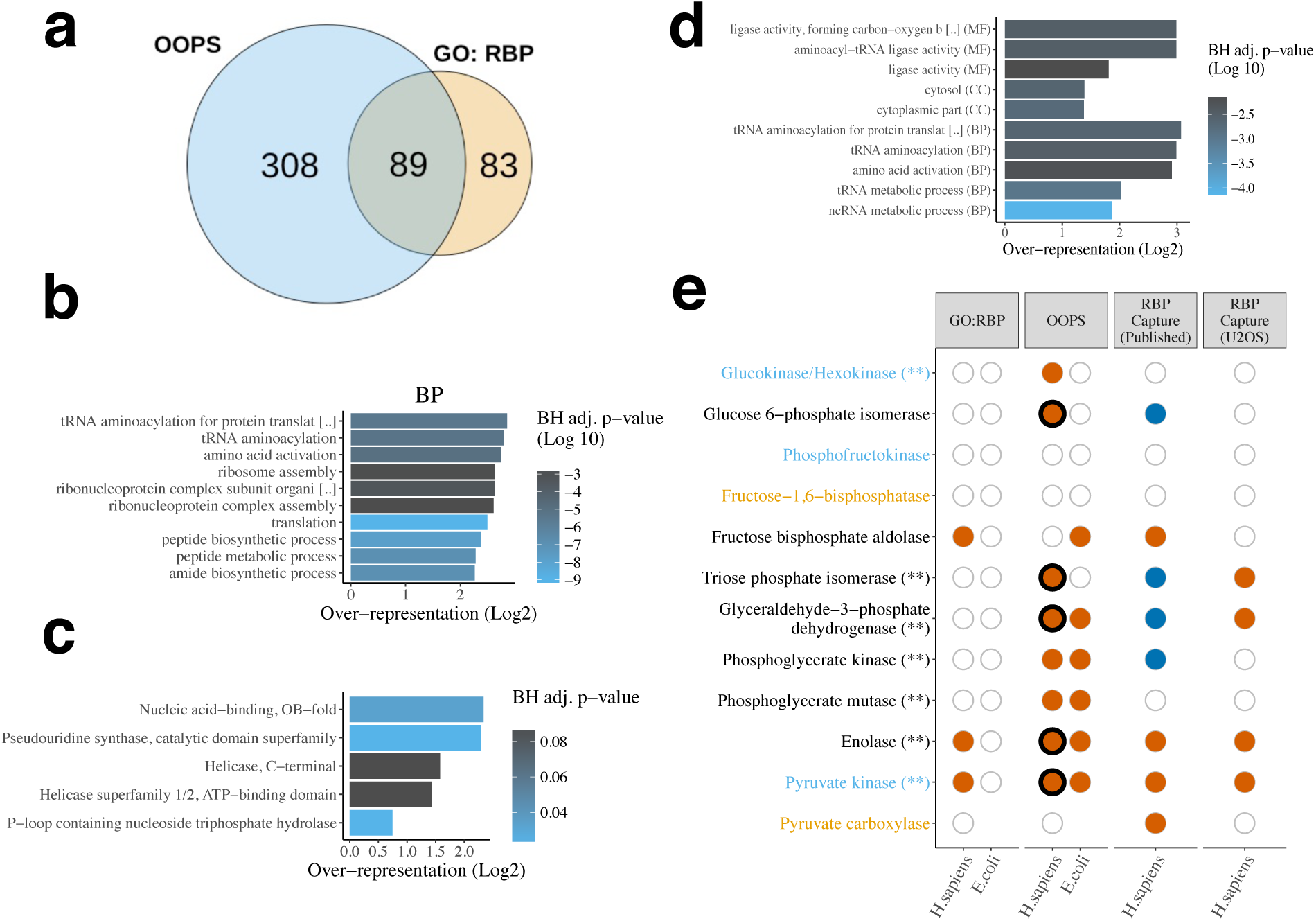
OOPS unravels the first bacterial RBPome. **(a)** Overlap between *E.coli* OOPS RBPs and GO annotated RBPs **(b)** Top 10 biological process GO terms over-represented in *E.coli* OOPS RBPs **(c)** Protein domains over-represented in *E.coli* OOPS RBPs **(d)** GO terms over-represented in OOPS novel *E.coli* RBPs **(e)** RNA-binding capacity of glycolysis/gluconeogenesis proteins. Proteins coloured by pathways; blue text = only glycolysis, orange text = only gluconeogenesis, black text = both, Asterisks = dynamic RNA-binding after nocodazole arrest. GO:RBP=GO-annotated RBP. Orange filled circle = protein observed in the dataset indicated. Dark blue fill = protein in human RBP-Capture experiments but listed as a lower-confidence “candidate” RBP. Empty circle = protein present in species but not observed in dataset. Thick black line indicates an RNA-binding site was identified in the sequential digestion experiment.

Interestingly, many of the glycolytic enzymes that bind RNA in *H. sapiens*, also bind RNA in *E. coli* (Figure 6e). Enolase 1 and Pyruvate kinase, detected by previous RBP-capture studies were identified as RBPs by OOPS in *E.coli*. Furthermore, GAPDH and PKG, previously described as low-confidence candidate RBPs in human by RBP-Capture, were also found as RBPs in our human and prokaryotic studies. Surprisingly, phosphoglycerate mutase, a glycolytic protein not previously identified in any human RBP-capture, was also detected as a RBP in both our eukaryotic and prokaryotic OOPS experiments. Taken together, the changes in RNA binding detected in our nocodazole experiment and the consistent identification of particular glycolytic RBPs between human and *E.coli* may indicate a conserved link between metabolism and RNA binding.

## Discussion

OOPS is able to retrieve crosslinked RNAs representing the complete cellular transcriptome and recover crosslinked RBPs, and strongly agrees with data obtained from other RBP identification methods. Importantly, our approach also unveils new RBPs from underrepresented subcellular compartments. We have further identified RNA-binding sites, providing independent evidence of specific RNA-protein interactions. Finally, we have assessed the ability of OOPS to analyse dynamic systems in a straightforward manner and its independence of requirement on a poly-A tail by describing the first bacterial RBPome.

Although OOPS recovers RNAs in an unbiased manner, we observe an under-representation of small RNAs (sRNAs) in the PBR fraction. We believe the most likely explanation is that small RNAs are less frequently protein-bound^37^. It is also possible that there is a lower probability of UV crosslinking of sRNAs to protein, as they have fewer RNA-protein interactions. Despite the apparent under-representation of sRNAs in our crosslinked interface, we consistently found sRNA-binding proteins in both human and bacteria. Indeed, in *E. coli,* OOPS recovers both canonical (Hfq) and recently discovered (ProQ) sRNA binding proteins^38^.

OOPS is dependent on separation of macromolecules by their physicochemical properties. Under such a separation, glycoproteins and RNA-protein adducts cannot be distinguished since both glycans and RNAs are hydrophilic polymers. Our observation that the interface abundance of the vast majority of the glycoproteins is CL-independent and RNase insensitive could indicate that they do not bind RNA. Alternatively, although very few glycoproteins have been described as RBPs, it is also possible they are true RBPs but their glycans maintain them in the interface even after RNase digestion. Despite this, it is interesting to note that 18/21 glycoproteins enriched by CL are localised to the exosome - a RNA-rich compartment ^39–41^ - and include 3 known RNA binding glycoproteins^10,42^. Yet, to enable a proper interrogation of RNA-binding glycoproteins, it would be necessary to remove glycans and achieving this in a manner that does not degrade RNA is non-trivial. We therefore took a conservative approach and discounted glycoproteins from our analyses.

Crosslinking-based detection of RBPs is based on proximity of RNAs and proteins. The functional relevance of some catalogs may therefore need to be considered with caution ^19^. Dynamic experiments, such as the one presented here, can facilitate interrogation of the biological function of the RNA-protein interactions and uncover system-wide changes in RNA-binding proteins. One of the most striking findings is the coordinated increase in RNA-binding of glycolytic enzymes following release from nocodazole arrest. Considering the previously described regulation of glycolytic protein thermal stability in response to nocodazole arrest^43^, and the reported repression of translation by GAPDH in response to changes in glycolytic flux^44^, our results suggest that an interplay between metabolism and RNA-binding may be a fundamental component of the cell cycle. GAPDH has been shown to bind to a range of RNA species including tRNAs, AU-rich elements, and TERC ^45,46^. *In vitro* competition assays and spectrometry models suggest binding occurs within its NAD-binding crevice, but this has not been observed *in vivo*^31,45,47^. Here, we provide the first *in vivo* evidence, which shows that the RNA-binding site of GAPDH is indeed the NAD-binding crevice.

In summary, OOPS is a low-cost method which allows the isolation of RNA-protein complexes in any organism, enabling downstream analysis of either the RNA or protein component. This simple method will make the study of RNA-protein interactions more accessible and foster a systems biology view of their function by permitting the study of their dynamic properties.

## Author Contributions

KSL, AW, EV, TS and RQ conceived the study. EV, RQ, TS and KSL designed the experiments. EV optimised the initial OOPS protocol, prepared RNA-seq libraries and performed flow cytometry analysis. EV performed the SILAC, cellular subfractionation and nocodazole arrest experiments with assistance from RQ. *E. coli* experiments were performed by MM, EV and RQ. U2OS RBP-Capture was performed by MP, VD. EV and RQ performed all additional experiments including the RNA binding site experiment. RQ performed all mass spectrometry. TS performed all data analysis, with the exception of the analysis of uridine content (DM) and analysis of *E. coli* data (TS, MM). EV, TS, RQ and KSL interpreted results, with critical appraisal of findings from AW, MP, MR, RH and VD, including additional experiments (MP). GT assisted with interpretation of *E. coli* data. MM-S performed the protein-RNA structural analysis. SD and LM developed the ionbot search engine and performed ionbot searches. TS, EV, RQ, KSL and MM-S drafted the manuscript, with critical revision from AW, RH, GT, DM, MP, MR and SD.

## Acknowledgments

EV, TS, RQ, RH, MP, MR, VD are supported by Medical Research Council, Grant/Award number: 5TR00; Wellcome Trust, Grant/Award numbers: 110170/Z/15/Z, 110071/Z/15/Z. MM-S is supported by the MRC (MC_U105185859) and by a FEBS Long-Term Fellowship. GT is supported by IB Catalyst grant for Project DETOX (BB/N01040X/1).

We would like to thank Harriet T. Parsons for donating *E. coli* cells, Mohammed A. Elzek for culturing MCF10A cells, Tom Mulroney for helping culturing U2OS cells and Bettina Fisher for kindly sharing equipment.

## Methods

### Cell culture

U-2 OS (U2OS) and MCF 10A cells were obtained from the American Type Culture Collection (ATCC). HEK-293 were kindly provided by Dr. Johanna Rees (University of Cambridge). U2OS and HEK-293 cells were cultured in McCoy’s 5A and DMEM (Gibco-BRL) media respectively, supplemented with 10% fetal bovine serum (Gibco-BRL). MCF 10A were maintained in MEBM media (Lonza/Clonetics) supplemented with 10 ng/ml of cholera toxin (Sigma-Aldrich). All cells were maintained at 37 °C and 5% CO2 and regularly tested for mycoplasma contamination with negative results.

E. coli K-12 DH5a strain (Thermo Fisher Scientific), was cultured in LB Broth (Thermo Fisher Scientific) at 37 °C. All E. coli experiments were done at stationary phase after 16 h of cell growth.

### Orthogonal Organic Phase Separation in human cells

Cells were cultured in 6 cm diameter dishes (28.2 cm^2^) for catalog experiments, or 10 cm diameter dishes (78.5 cm^2^) for dynamic experiments, until a maximum of 90% of confluence was reached, using a single dish per replica and condition. Cells were washed twice with PBS and supernatant removed by pipetting. In non-crosslinked controls, cells were immediately lysed by scrapping in Acidic Guanidinium Thiocyanate-Phenol (Trizol, Thermo Fisher Scientific), and the homogenate transferred to a new tube. In crosslinked samples, UV-crosslinking was performed on PBS-washed cells by UV-irradiation at 254 nm (CL-1000 Ultraviolet Crosslinker; UVP). Immediately after crosslinking, cells were scraped in Trizol and the homogenized lysate was transferred to a new tube and incubated at room temperature (RT) for 5 min to dissociate unstabilised RNA-protein interactions. For biphasic extraction, 200 μL of chloroform (Fisher Scientific) were added, phases were vortexed and centrifuged for 15 min at 12,000 ×; *g* at 4 ^°^C. The upper aqueous phase (containing non-crosslinked RNAs) was transferred to a new tube, and RNA precipitated following manufacturer instructions. The lower organic phase (containing non-crosslinked proteins) was transferred to a new tube and proteins precipitated by addition of 9 volumes of methanol (Fisher Scientific). Interface (containing the Protein-RNA adducts) was subjected to extra AGPC phase separation cycles, precipitated by addition of 9 volumes of methanol, and pelleted by centrifugation at 14,000 x g, RT for 10 min.

For RNA analyses, the precipitated interfaces were incubated for 2h at 50 °C in 30 mM Tris HCl (pH8)/10 mM EDTA and 18 Us of proteinase K (Thermo Fisher Scientific). Samples were cooled and released RNA was purified by standard phenol/chloroform extraction (Thermo Fisher Scientific) according to the manufacturer instructions.

For RNA-binding protein analyses, the precipitated interface was resuspended in 100 μL of 100 mM TEAB, 1 mM MgCl_2_, 1% SDS, incubated at 95 °C for 20 min, cooled down and digested with 2 μg RNAse A, T1 mix (2 mg/mL of RNase A and 5000 U/mL of RNase T1, Thermo Fisher Scientific) for 2-3 h at 37 °C. Another 2 μg of RNAse mix was added and incubated overnight at 37 °C, after which a final cycle of AGPC phase partitioning was performed and released proteins recovered from the organic phase by methanol precipitation.

### Orthogonal Organic Phase Separation in bacteria

*E. coli* cultures were grown overnight. 3 ml of culture was pelleted by centrifugation (5 min at 6000 g, RT) and washed twice with PBS. Cells were resuspended in PBS and crosslinked in solution at 254 nm for 525 mJ/cm^2^. Crosslinked cells were pelleted again and supernatant removed by pipetting, leaving approximately 50 μl of PBS. 500 μl of 0.5 mm glass beads (Sigma-Aldrich) were added to each sample, mixed gently, frozen on dry ice and dried by sublimation for 2 h. Dried cells were disrupted by vortexing for 5 min, at intervals of 1 min to avoid warming the sample. 1 ml Trizol was added to each tube and samples were homogenized by vortexing. Supernatant (avoiding glass beads) was transferred to a new tube and centrifuged 5 min at 6000 x g at 4 °C. The supernatant was transferred to a new tube, leaving the unlysed cells as a pellet. Finally, OOPS was performed as described above.

### RNA quantification and integrity assessment

RNA purity was assessed by Nanodrop (Thermo Fisher Scientific). Samples with a 260/280 ratio below 1.9 or 260/230 below 2 were discarded. RNA concentration was estimated using the Qubit RNA BR (Broad-Range) Assay Kit (Thermo Fisher Scientific) in the Qubit^®^ 2.0 Fluorometer (Thermo Fisher Scientific). RNA integrity was evaluated using the Agilent 2100 Bioanalyzer system (Agilent).

### RNA sequencing

Protein Bound RNA (PBR) and total non-crosslinked (NC) RNA were purified using OOPS or standard Trizol extraction respectively. All RNA samples were treated with turbo DNase (Thermo Fisher Scientific). Ribosomal RNA (rRNA) was depleted using RiboCop kit V1.2 (Lexogen, Greenland, NH, USA) according to manufacturer instructions, starting with 1ug of RNA. Two nanograms of rRNA-depleted NC-RNA or 8 ng of rRNA-depleted PBR were used to generate sequencing libraries using SENSE total RNA-Seq Library Prep kit (Lexogen). All libraries were sequenced in parallel on a NextSeq 500 for 75 cycles (Illumina).

### RNA-Seq data processing and bioinformatics

Quality control of raw fastqs was performed using FastQC (www.bioinformatics.babraham.ac.uk/projects/fastqc/). Reads were aligned to the hg38 human genome and Ensembl 87^48^ using hisat2^49^ with default settings and reads with MAPQ < 10 were discarded. Transcript quantification was performed with Salmon^50^ using default settings. The meta-plot of read coverage over gene model was obtained using the CGAT bam2geneprofile script with *reporter=utrprofile*^51^. For details of the identification of putative RNA binding sites and the overlap with eCLIP data, see supplementary notes.

### Oligo(dT) RBP-capture

RBP-Capture was performed according to^23^, with the following modifications. We used 4 ×; 500cm^2^ plates per condition. Oligo(dT)25 magnetic beads (NE Biolabs) were reconditioned as per manufacturer’s instructions and incubated with the lysates for a second round of RBP-capture with eluates from the two rounds were pooled together

### Subcellular fractionation

U2OS cells from a single 80% confluent 500 cm^2^ cell culture dish (Sigma-Aldrich) were detached using trypsin without EDTA (Thermo Fisher Scientific), pelleted 5 min at 250 g, washed with PBS, resuspended in 50 ml of PBS and crosslinked in solution at 254 nm at 400 mJ/cm^2^. Cells were pelleted again for 5 min at 250 x g, resuspended in 1 ml of lysis buffer (0.25 M sucrose, 10 mM HEPES pH 7.4) containing protease inhibitors (Roche), and lysed with a ball-bearing homogenizer (Isobiotec) on ice. Unlysed cells were removed by centrifugation at 200 x g, 5 min at 4 °C. The supernatant was transferred to a new tube and centrifuged at 1000 x g, 10 min at 4 °C with the pellet collected as ‘heavy membrane fraction’. The supernatant was centrifuged again at 12.200 g with the pellet collected as the ‘light membrane fraction’. The supernatant was collected as cytosolic fraction, frozen and dried by sublimation by SpeedVac (Labconco). Pellets from the heavy membranes, light membranes and cytosol were re-suspended in Trizol and RBPome and “total” proteome were extracted using OOPS.

### Nocodazole synchronisation

A single 10 cm^2^ diameter dish (per replica and condition) of U2OS cells at 70 % of confluence was arrested in prometaphase by direct addition of 1μg/ml of nocodazole (Sigma-Aldrich) to the cell culture media. 16-18 hours post treatment, synchronised cells were washed twice in PBS and crosslinked at 254 nm at 400 mJ/cm^2^. Arrested cells were detached by mechanical stimulation, pelleted, solubilised in Acidic Guanidinium Thiocyanate-Phenol (Trizol) and stored at −80 °C. For post-release timpoints (6 h and 23 h post-arrest), synchronised cells were detached form the dish by mechanical stimulation, washed in PBS and re-seeded in media without nocodazole. Cells were then washed twice with PBS and crosslinked at 254 nm at 400 mJ/cm^2^. Cell lysates were obtained by directly scraping the crosslinked cells in Acidic Guanidinium Thiocyanate-Phenol (Trizol). The total proteome was extracted from the lysate and the RBPome was determined using OOPS (see Orthogonal Organic Phase Separation in human cells). A parallel cell dish was cultured for every time point and replicate to assess the synchronisation efficacy by flow cytometry. DNA content per cell was analysed using the Propidium Iodide Flow Cytometry Kit (Abcam) as indicated by the manufacturer. Flow cytometry results were analysed using FlowJo 8.7.

### Proteomic sample preparation

Samples were resuspended in 100 μL of 100 mM Triethylammonium bicarbonate (TEAB) (Sigma-Aldrich), reduced with 20 mM DTT (Sigma-Aldrich) at room temperature for 60 min and alkylated with 40 mM iodoacetamide (Sigma-Aldrich) at room temperature in the dark for at least 60 min. Samples were digested overnight at 37 °C with 1 μg of Trypsin (Promega) with the exception of samples for TMT labeling which were digested overnight at 37 °C with 1 μg Lys-C (Promega). Subsequently, 1 μg of modified trypsin (Promega) was added, and the samples were incubated for 3-4 h at 37 °C. Samples were then acidified with TFA (0.1% (v/v) final concentration; Sigma-Aldrich) and centrifuged at 21,000 g for 10 min, with the supernatant frozen at −80 °C until required.

For peptide clean-up and quantification, 200 μL of Poros Oligo R3 (Thermo Fisher Scientific) resin slurry (approximately 150 μL resin) was packed into Pierce™ Centrifuge Columns (Thermo Fisher Scientific) and equilibrated with 0.1% TFA. Samples were loaded, washed twice with 200 μL 0.1% TFA and eluted with 300 μL 70% acetonitrile (ACN) (adapted from^52^). 10 μL was taken from each elution for Qubit^™^ protein assay (Thermo Fisher Scientific) quantitation, with the remaining sample retained for MS.

### LC-MS/MS

SILAC labelling was performed according to the manufacturer’s instructions by growing cells in DMEM media containing light (Arg0-Lys0) or heavy (Arg10-Lys8) isotopes (SILAC Protein Quantitation Kit, Thermo Fisher Scientific). SILAC and unlabeled samples generated from OOPS experiments were analysed in the Orbitrap Fusion Lumos (Thermo Fisher Scientific) coupled to a nanoLC Dionex Ultimate 3000 UHPLC (Thermo Fisher Scientific). Mass spectra were acquired using CHarge Ordered Parallel Ion aNalysis (CHOPIN) acquisition in positive ion mode as previously reported^53^

TMT-10plex (Thermo Fisher Scientific) labelling from desalted peptides was performed according to the manufacturer’s protocol. Equal amounts of desalted peptides were labelled immediately after being quantified with Qubit^™^ protein assay (Thermo Fisher Scientific). Multiplexed TMT samples were separated into 4 fractions using Pierce™ High pH Reversed-Phase Peptide Fractionation Kit (Thermo Fisher Scientific). TMT-10plex fractions were analysed in an Orbitrap Fusion Lumos. Mass spectra were acquired in positive ion mode applying data acquisition using synchronous precursor selection MS^3^ (SPS-MS^3^) acquisition mode^54^.

Samples from Oligo(dT) capture and from subcellular fractionation were analysed in an Orbitrap nano-ESI Q-Exactive mass spectrometer (Thermo Fisher Scientific), coupled to a nanoLC (Dionex Ultimate 3000 UHPLC).

### Peptide spectrum matching

Raw data were viewed in Xcalibur v.2.1 (Thermo Fisher Scientific), and data processing was performed using Proteome Discoverer v2.1 (Thermo Fisher Scientific). The Raw files were submitted to a database search using Proteome Discoverer with Mascot, SequestHF and MS Amanda algorithm^55^ against the *Homo sapiens* database for U2OS and MCF 10A cells or *E. coli* database, downloaded in early 2017 containing human (or *E. coli*) protein sequences from UniProt/Swiss-Prot and UniProt/ TrEMBL. Common contaminant proteins (several types of human keratins, BSA, and porcine trypsin) were added to the database, and all contaminant proteins identified were removed from the result lists prior to further analysis. The spectra identification was performed with the following parameters: MS accuracy, 10 ppm; MS/MS accuracy, 0.5 Da for samples run with Orbitrap Fusion Lumos and MS accuracy, 10 ppm; MS/MS accuracy, up to 0.2 Da for samples run with Q-Exactive mass spectrometer; up to two missed cleavage sites allowed; carbamidomethylation of cysteine (as well as TMT6plex tagging of lysine and peptide N-terminus for TMT labeled samples) as a fixed modification; and oxidation of methionine and deamidated asparagine and glutamine as variable modifications. Percolator node was used for false discovery rate estimation and only rank 1 peptide identifications of high confidence (FDR < 1 %) were accepted. A minimum of two high confidence peptides per protein was required for identification using Proteome Discoverer. TiO_2_-enriched fractions intended for direct assessment of RNA-crosslinked site in proteins were searched with beta-testing version *Ionbot* search engine (see Supplementary notes) with carbamidomethylation of cysteine and oxidation of methionine respectively as a fixed and variable modifications, *Homo sapiens* Swissprot database and both *Extended Modification Search* and *RNA Binding Modification Search* enabled.

### Direct assessment of RNA crosslinking site in proteins

Starting from the precipitated OOPS interface, proteins were digested using 1μg Lys-C (Promega, Madison, WI, USA) in 100 μL of 100 mM TEAB (Sigma-Aldrich) with 1 μL of RNAseOUT (Thermo Fisher Scientific) overnight at 37 °C. Two different approaches were used to enriched RNA-peptides:

i. Silica-based RNA purification using the RNeasy kit (Qiagen), according with the manufacturer’s instructions;
ii. Precipitation in 80% ethanol. Two rounds of precipitations were used to further clean the sample.

RNA-peptides were re-suspended in 100 μL of 100mM Tris-HCl (pH8.0)/ 2 mM MgCl_2_, sonicated for 15 min and incubated at 95 °C for 20 min. 2 μg RNAse A/T1 mix (2 mg/mL of RNase A and 5000 U/mL of RNase T1) was added to cooled samples, and incubated for 4 h at 37 °C followed by a second protease digestion using 1 μg trypsin (Promega) overnight at 37 °C. Digested samples were desalted with Oligo R3 as described in the “proteomics sample preparation” section and dried on speedvac (Labconco).

Digests were re-suspended in 30-40 μL of 80% acetonitrile (ACN)/2% TFA containing 1 μg of TiO_2_ beads (GL Sciences). The slurry was transferred into a p200 tip containing a C8 “plug” (3M Empore, Sigma-Aldrich) to retain the loaded TiO_2_ beads and the flow-through collected. The packed TiO_2_ was washed with 20 μL 80% ACN/2% TFA, then 20 μL 10% ACN/0.1% TFA and the flow-through from both retained. The TiO_2_-enriched fraction was eluted from the beads with two rounds of 20 μL of Ammonia solution (1.5-1.8%), pH>10.5, and 20 μL of 50% ACN.

### Proteomics bioinformatics and data analysis

Peptide-level output from Proteome Discoverer was re-processed with the add_master_protein.py script (https://github.com/TomSmithCGAT/CamProt) to ensure uniform peptide to protein assignment for all samples from a single experiment and identify peptides which are likely to originate from contaminating proteins such as keratin (see supplementary notes). For quantitative experiments, peptide-level quantification was obtained by summing the quantification values for all peptides with the same sequence but different modifications. Protein-level quantification was then obtained by taking the median peptide abundance. For SILAC experiments, the ratio between treatment and control protein abundance was calculated for each sample separately and aggregated to average protein ratio. For TMT experiments, data analysis was performed using the *MSnbase* R package^56^. Log_2_-transformed protein abundance was centre-median normalised within each sample. For the crude fractionation experiments (*n*=5), the protein abundance was quantified by label-free quantification, averaged across the replicates per fraction and normalised per protein such that the sum of abundances over the 3 fractions was 1. For the U2OS RBP-Capture experiment, only proteins observed in all 3 CL replicates and no NC replicates were retained. In crosslink-testing SILAC experiments, only proteins present in at least 2 replicates were retained.

GO terms, Interpro protein domains and KEGG pathway annotations were obtained using the R package *UniProt.ws*^57^. GO terms were expanded to include all parent terms using the R package *GO.db*^58^. Glycoproteins were identified using the Uniprot^59^ API with *categories=PTM* and *types=CARBOHYD*. Transmembrane proteins were identified using the Uniprot API with *types=TRANSMEM*.

### Data handling and statistics

Data handling was performed with R v3.4.1 using tidyverse packages and python v3.6.5. Plotting was performed with the *ggplot2* R package^60^.

Proteins observed only in CL in at least one replicate were deemed enriched. For the RNAse-testing SILAC experiments, proteins only ever observed in the RNAse condition at the organic phase were deemed enriched. Vis versa, those only ever observed in the control condition at the interfaces were deemed depleted. For proteins which did not meet these criteria, all peptides observed across the replicates were treated as independent observations and a Mann-Whitney-Wilcoxon Test was used to test whether the log_2_ median CL:NC or RNase:Control ratio was > 0 (enriched) or < 0 (depleted), with a BH-adjusted p-values < 0.05 considered significant. Proteins with less than 6 peptides were excluded from the statistical test due to insufficient power.

GO, InterPro and KEGG over-representation analyses were conducted using the R package goseq to account for protein abundance. p-values were adjusted to account for multiple testing using the Benjamini-Hochberg^61^ FDR procedure. GO-terms with adjusted p-value <0.01 and at least 5 proteins were considered significantly over-represented. Over-representation values given are not adjusted for protein abundance. For U2OS and HEK-293, protein abundance was derived from^62^ taking the maximum abundance recorded across the replicates. For MCF10A, we used an in-house deep proteomics data set. For *E.coli*, protein abundance was obtained from PaxDB^63^.

For the nocodazole arrest/release experiment, proteins with a change in abundance or RNA binding were identified using the *lm* function in R. Specifically, to identify protein with a change in abundance between nocodazole arrest and 6 h release, total protein abundance was modelled as a function of the time point alone (abundance ∼ timepoint). The p-values for the timepoint coefficients for each proteins were adjusted to account for multiple hypothesis testing according to Benjamini-Hochberg^61^ and proteins with an adjusted p-value < 0.01 (1 % FDR) were considered to have changed abundance. To identify proteins with a change in RNA binding between nocodazole arrest and 6 h release, protein abundance in the total proteome and OOPS samples was modelled as a function of the time point, the abundance type (total or OOPS), and the interaction between these two variables (abundance ∼ timepoint + type + timepoint*type). Here, the interaction term denotes whether the abundance in OOPS and total follows the same pattern across the timepoints (coefficient is zero), indicating total abundance determines the amount of protein bound to RNA, or diverges (non-zero coefficient), indicating a change in RNA binding between the timepoints. The p-values for the interaction term were obtained and adjusted as indicated above. For the heatmap representation, protein abundances were z-score normalised within the total and OOPS samples separately. Hierarchical clustering was performed with the R *hclust* function using 1-Spearman’s rho as the distance metric and average linkage.

For details of the identification of RNA binding sites see supplementary notes.

### Structural Assessment of RNA-protein contacts

In order to look for structural information to validate our direct evidence for RNA-protein contacts, the Uniprot IDs of the detected proteins were used to retrieve all their associated PDB IDs using the Uniprot Retrieve/ID mapping tool. In parallel, we retrieved information for all structures annotated as containing protein-RNA complexes in the nucleic acid database^64^. Comparison of PDB IDs common in both subsets revealed the structures of the ribosome quality control complex (PDB ID 3J92) and of a Glycyl-tRNA synthetase in complex with tRNA-Gly (PDB ID 4KR2). These structures, together with the structure of GADPH in complex with NAD (PDB ID 4WNC), were later visualized using VMD 1.9.4^65^.

